# Lung Microbiome Intervention Attenuates Herpesvirus-Induced Post-HCT Pulmonary Fibrosis Through PD-L1 Upregulation on Dendritic Cells

**DOI:** 10.1101/2024.07.06.602351

**Authors:** Joshua B. Perkins, Keerthikka Ravi, Chunfang Guo, Gina J. Oh, Bellmary G. Rodriguez, Stephen J. Gurczynski, Jason B. Weinberg, Gary B. Huffnagle, David N. O’Dwyer, Bethany B. Moore, Xiaofeng Zhou

## Abstract

Alterations in the lung microbiome frequently accompany adverse pulmonary outcomes. Hematopoietic cell transplantation (HCT) markedly affects the lung microbiome corresponding with a high incidence of post-HCT pulmonary complications. In a preclinical mouse model of HCT, we observed a reduction in *Lactobacillus johnsonii* within the lung microbiome following transplantation. Intranasal administration of live or heat-killed (HK) *L. johnsonii* at low doses reduced gammaherpesvirus-induced pulmonary fibrosis in HCT mice, in which IL-17A plays an essential role. HK *L. johnsonii* treatment of HCT mice suppressed inflammatory cytokine production by lung macrophages and decreased *Il17a* expression in T helper 17 (Th17) cells. HK *L. johnsonii* increased PD-L1 expression on the surface of type II conventional dendritic cells (cDC2) in HCT mice and *in vitro* in bone marrow-derived dendritic cells (BMDCs). HK *L. johnsonii*-exposed BMDCs also inhibited IL-17A secretion from co-cultured Th17 cells in a PD-1-dependent manner. Notably, when HK *L. johnsonii* was administered to HCT mice reconstituted with bone marrow cells from PD-1 knockout (KO) mice, which lack a PD-L1 mediated response, HK *L. johnsonii*-mediated reduction of pulmonary fibrosis was negated. Collectively, our findings demonstrate that HK *L. johnsonii* mitigates herpesvirus-induced pulmonary fibrosis in HCT mice by modulating cDC2 surface expression of PD-L1, which subsequently suppresses *Il17a* expression in Th17 cells, pointing towards a potential postbiotic-based strategy for immunomodulation to address pulmonary complications of HCT.

## Introduction

The lung microbiome is characterized by low biomass and a constant state of flux; however, various features of the bacterial communities in the lung microbiome have been found to correlate with pulmonary pathology^1^. For instance, in patients with idiopathic pulmonary fibrosis (IPF), increased bacterial burden and alterations in the microbial community structure, combined with a reduction in diversity, are linked with the progression of the disease^2,3^. As our understanding of the lung microbiome advances, the possibility of developing microbiome-based therapeutics aimed at restoring the function of a normal healthy lung microbiome emerges. However, there are significant challenges to overcome before these can be applied clinically. These include the identification of precise microbial targets for specific lung conditions, elucidating the mechanisms by which these targets influence lung health, and determining the most effective formulations and routes for therapeutic delivery^1,4,5^.

Medical procedures frequently disrupt the host’s microbiome, which in turn can be linked to adverse clinical outcomes. This is particularly evident in the context of hematopoietic cell transplantation (HCT), where both the procedure itself and the multiple exposures to antibiotic treatment that are common in HCT patients can substantially alter the gut microbiome^6^ and the lung microbiome^7^. Pulmonary complications, affecting up to 60% of allogeneic HCT recipients^8^ and 25% of autologous HCT recipients^9^, are often preceded by dysbiosis in the lung and gut microbiome, suggesting a predictive relationship between microbial imbalances and respiratory challenges following transplantation^10^.

Idiopathic pneumonia syndrome (IPS) is a serious non-infectious pulmonary complication of HCT that is characterized by pneumonitis, lung injury, and often fibrosis, leading to irreversible morbidity and mortality^11^. The specific causative factors of IPS are not completely defined. However, we previously demonstrated a marked association between herpesvirus infection shortly after transplant, such as human herpesvirus 6 (HHV-6) and Epstein-Barr virus (EBV), and IPS development in human HCT recipients^12^. In preclinical studies, we found that acute infection with murine gammaherpesvirus 68 (MHV-68), a mouse virus with similarities to the human gammaherpesvirus EBV, following syngeneic (syn) HCT in mice, induces an IL-17-dependent inflammatory pneumonitis and subsequent progression to fibrotic lung disease, mirroring key elements of human IPS pathology^13–15^. Intriguingly, a notable dysbiosis occurs within the lung bacterial community post-HCT, particularly a significant depletion of the *Lactobacillus* genus^7^. In bronchoalveolar lavage (BAL) fluid from human HCT recipients diagnosed with IPS, the *Lactobacillaceae* family members were nearly undetectable^7^. Many *Lactobacillus* species/strains are recognized as probiotics and have the potential to modulate specific mucosal immune system functions^16^. However, the role of lactobacilli in the development of pneumonitis and pulmonary fibrosis following HCT and the underlying mechanisms of their action have not yet been investigated.

In this study, we discovered that the strain *Lactobacillus johnsonii* XZ17 (GenBank accession number CP151183.1) is prevalent in the lungs of healthy non-HCT mice but is diminished following HCT. We found that intranasal administration of either live or heat-killed (HK) *L. johnsonii* XZ17 to HCT mice mitigates pneumonitis and pulmonary fibrosis. Treatment with HK *L. johnsonii* XZ17 increased PD-L1 expression on type II conventional dendritic cells (cDC2) and PD-1 on T helper 17 (Th17) cells, which in turn suppressed the production of IL-17A, a key mediator of pneumonitis and fibrosis in HCT mice. Additionally, HK *L. johnsonii* XZ17 exerted anti-inflammatory effects by inhibiting the expression of pro-inflammatory cytokines such as *Il6*, *Il1b*, *Il18*, and *Tgfb1* in lung macrophages. The use of heat-inactivated *L. johnsonii* offers enhanced safety for immunotherapy in immunocompromised patients. Overall, our findings have identified a specific microbial target that could be modulated and acts in a PD-L1/PD-1 dependent manner to potentially offer a safe and effective strategy to reduce pulmonary complications post-HCT.

## Results

### *Lactobacillus johnsonii* in the mouse lung microbiome declined following syngeneic HCT

HCT itself, as well as the subsequent medications and infectious complications, profoundly alter the lung microbiome^7,17^. We previously developed a mouse model in which syn-HCT followed by MHV-68 infection led to severe pneumonitis and pulmonary fibrosis^13,15^. We have shown that, similar to its effects in humans, HCT modifies the lung microbiome composition in mice, and a subsequent MHV-68 infection further reduces community diversity^7^. We also found that the relative abundance of members of the genus *Lactobacillus* was significantly reduced in the lungs of mice following HCT^7^. To verify these results, we cultured lung homogenates from non-HCT or HCT mice on de Man–Rogosa–Sharpe (MRS) agar, a selective medium that promotes the growth of lactobacilli^18^. Lungs from non-HCT mice harbored a higher number of culturable lactobacilli compared to those from HCT-treated mice (**Fig. 1a**), Among those tested lungs, 10 out of 15 (66.7%) of the non-HCT mouse lung samples yielded lactobacillus colonies on MRS agar, whereas only 4 out of 15 (26.7%) of the samples from HCT recipients did so (**Fig. 1b**). These colonies were morphologically consistent, composed of Gram-positive rods (**Fig. 1c**). Eight colonies were randomly selected for molecular analysis. We performed bidirectional sequencing of the 16S rRNA gene using the D88 forward and E94 reverse degenerate primers^19^. The sequencing resulted in high-quality reads with an approximate length of 1170 base pairs, revealing that the sequences of all isolates were identical. The sequences showed complete identity with the known 16S rRNA gene sequence of *Lactobacillus johnsonii*^20^. Thus, we have identified a bacterial species that is significantly diminished in the lungs after HCT in mice.

**Figure 1.**
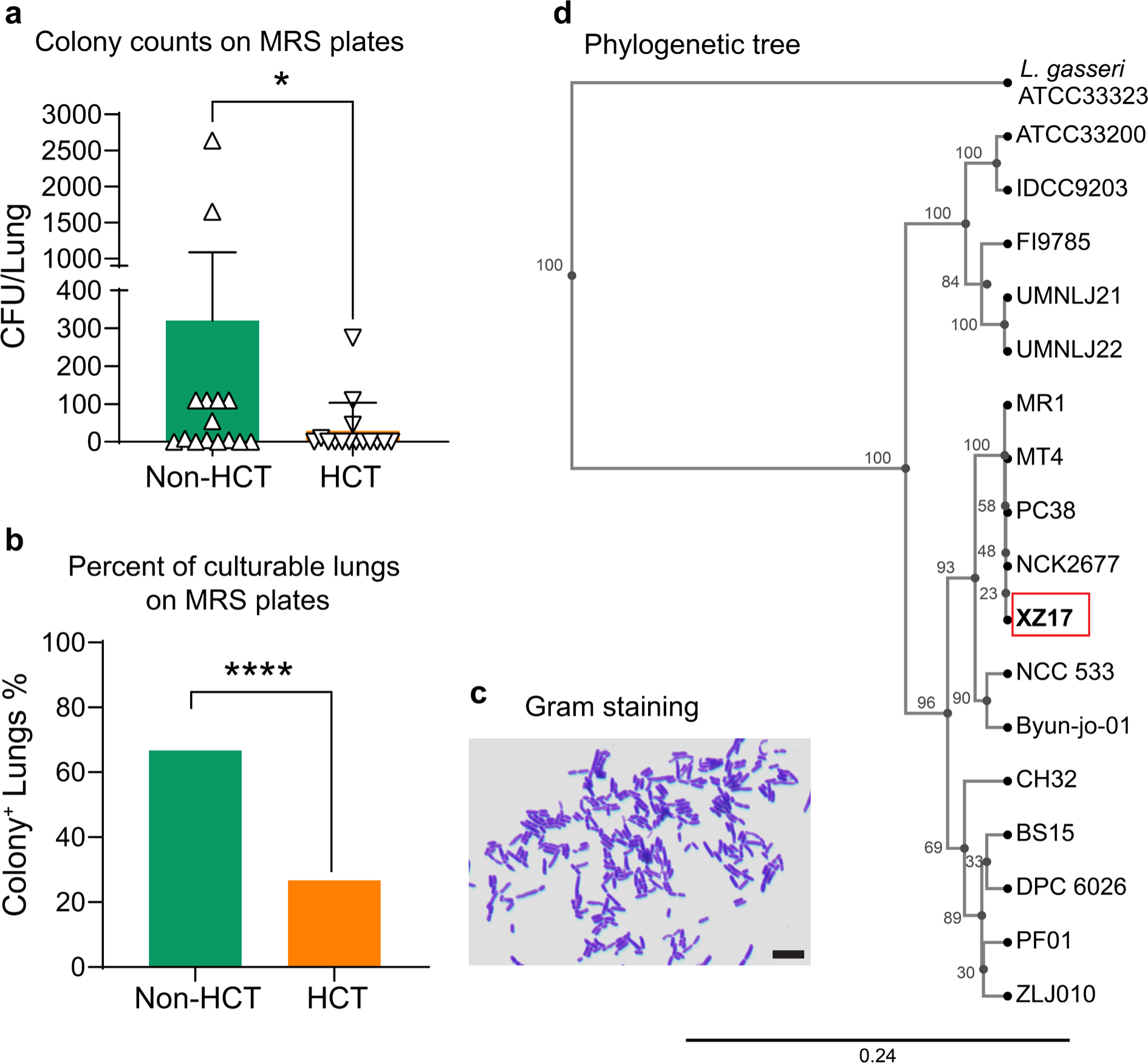
*Lactobacillus johnsonii* is diminished following syngeneic HCT in mouse lungs. **a**. Lungs were isolated from 15 C57BL/6J mice five weeks after receiving synthetic HCT or from age-matched, healthy non-HCT controls. Each lung was homogenized and subsequently plated onto MRS agar plates for the quantification of *Lactobacillus* colony-forming units (CFUs). The CFUs per lung homogenate were counted 24 hours after plating. **b**. Percentage of mouse lungs with positive *Lactobacillus* cultures (n=15 per group). **c**. Gram staining of bacterial colonies cultured from the lungs of a non-HCT C57BL/6J mouse. Scale bar, 5 μm. **d**. Pairwise whole genome sequence average nucleotide identity (ANI) score-based phylogenetic tree showing the relationship of the *L. johnsonii* XZ17 strain (red rectangle) to 17 other *L. johnsonii* sequences sourced from the NCBI database, along with the *L. johnsonii* type strain ATCC33200. *L. gasseri* ATCC33323, a closely related species is used as the out-group for the analysis. The phylogenetic tree was built using hierarchal clustering of the Euclidean distance matrix of pairwise ANI scores. The number on the nodes represents the bootstrap value. The bar at the bottom provides the scale of the branch length. For Fig. 1a, data are expressed as mean ± SEM, with \**P*<0.05 as determined by the Mann-Whitney U test. For Fig. 1b, \*\*\*\**P*<0.0001 was determined by Fisher’s exact test.

We further sequenced the whole genome of a selected colony, designated as *L. johnsonii* strain XZ17, and submitted the sequence to GenBank (accession number CP151183.1) as detailed by Ravi et al. (2024, submitted). The genome of strain XZ17 comprises 1,948,040 base pairs with a GC content of 34.64%, and it contains 1,780 predicted protein-coding genes. Phylogenetic analysis revealed that the genome of XZ17 shares a high degree of similarity with other mouse isolates from the Jackson Laboratory, including MR1^21^, MT4^22^, PC38^23^ and NCK2677^24^, exhibiting an average nucleotide identity (ANI) of 99.9%. XZ17 also has a significant similarity of 98.4% with the human probiotic isolate NCC 533^25^ which clusters with all mouse isolates in the phylogenetic tree (**Fig. 1d**). Notably, the related MR1 strain, which becomes enriched in the cecum of BALB/cJ mice after administration with house dust collected from homes with indoor/outdoor dogs, has been shown to enhance airway immune defense against allergens and respiratory syncytial virus (RSV) infection when administered intragastrically^26,27^.

### Intranasal administration with either live or nonviable *L. johnsonii* XZ17 mitigates pulmonary fibrosis in HCT mice following MHV-68 infection

*L. johnsonii* can reduce inflammation and mucogenic responses caused by RSV infection^26–28^, and as shown above, it is significantly diminished after HCT (**Fig. 1b** and **c**). Based on this evidence, we chose to reintroduce the *L. johnsonii* XZ17 strain into the lungs of HCT mice. To determine the colonization potential of the XZ17 strain in the lungs of HCT mice, 5×10^5^ colony-forming units (CFU) XZ17 were administered intranasally to HCT mice. This low dose is presumed to be physiologically relevant, as it mirrors the typically low biomass of the lung microbiome compared to that of the gut microbiome^29^. Our results indicated an approximately two-log reduction of XZ17 CFUs one day post-instillation, and by day three, virtually no culturable XZ17 remained (**Supplementary Fig. 1a**). Live/dead bacterial staining (**Supplementary Fig. 1b**) revealed that despite the absence of culturable XZ17 in the lungs three days after instillation, a small proportion of the XZ17 bacteria remained in a viable but nonculturable state (**Supplementary Fig. 1c**). These results imply that the HCT lung environment is inhospitable for *L. johnsonii* colonization, likely owing to conditions such as elevated oxidative stress and insufficient nutrient availability. This result also suggests that lactobacilli normally found in the lungs of healthy mice are likely transient visitors, routinely dispersed from the upper respiratory tract, rather than permanent inhabitants.

**Supplementary Figure 1:**
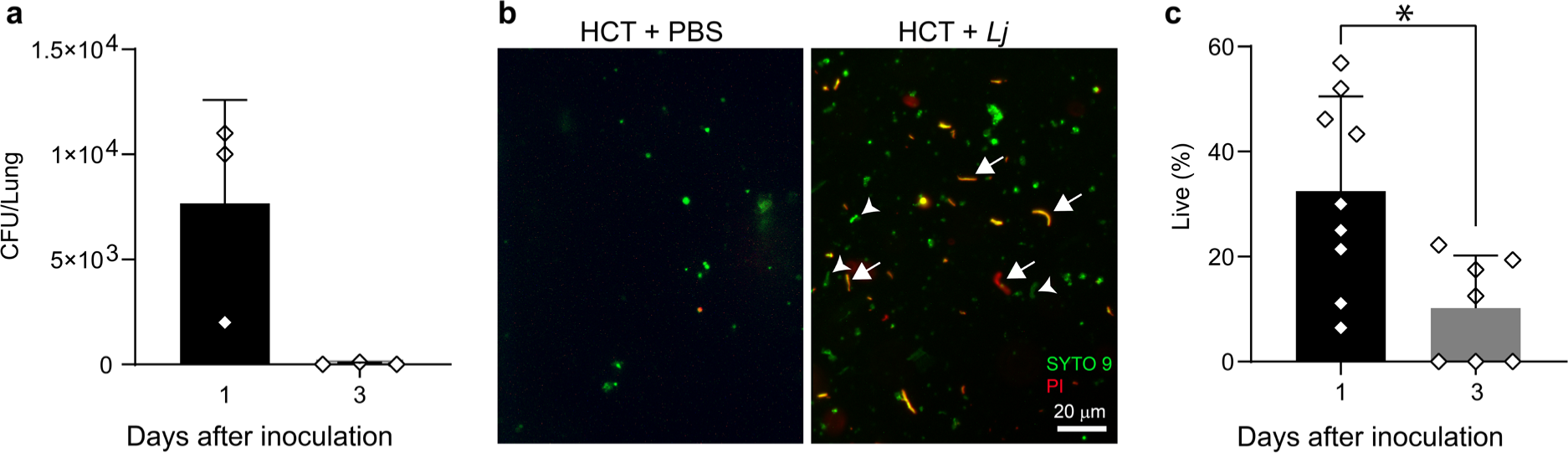
Intranasally inoculated *L. johnsonii* XZ17 fails to establish colonization in the lungs of HCT mice. **a**. Colony-forming units (CFU) assessed by culturing lung homogenates on MRS agar one or three days post-inoculation with 5×10^5^ CFU of *L. johnsonii* XZ17. **b**. Representative image displaying live/dead staining of lung isolates from HCT mice three days after inoculation with *L. johnsonii* XZ17 (HCT+*Lj*) or vehicle (HCT+PBS). Arrowheads highlight live rod-shaped bacteria stained green; arrows indicate dead or dying bacteria stained red or yellow, respectively. **c**. Percentage of live rod-shaped bacteria in lung isolates from HCT mice one or three days after inoculation with *L. johnsonii* XZ17. Data are presented as mean ± SEM. \**P*<0.05, determined by unpaired two-tailed Student’s *t* tests. Results shown are representative of two independent experiments.

To assess the impact of XZ17 on the lungs of HCT mice, we administered 5×10^5^ CFU of live XZ17 or an equivalent quantity of heat-killed (HK) XZ17 intranasally to HCT mice every 2 or 3 days, beginning 7 days prior to MHV-68 infection and continuing for a duration of 3 weeks post-infection (**Fig. 2a**). HK XZ17 was prepared by incubating the bacteria in a 70°C water bath for 15 minutes. For the oral route, we administered live 4×10^7^ CFU XZ17 via oral gavage following the same schedule as intranasal administration. Previous studies have demonstrated that oral administration at this dosage can alleviate pathology associated with RSV infection^26,27^. Results showed that intranasal administration of either live or HK XZ17 significantly reduced collagen content and collagen type I alpha 1 chain (*Col1a1*) expression in the lungs of HCT mice at 21 days post-infection (dpi) with MHV-68, as compared to levels measured in mock-inoculated controls (**Fig. 2b**). In contrast, high-dose oral administration of live XZ17 did not significantly attenuate pulmonary fibrosis in MHV-68-infected HCT mice (**Fig. 2b**). Histological examination of the lungs using Masson’s Trichrome staining supported the differential impact of intranasal versus oral administration on collagen accumulation (**Fig. 2c** and **d**). The fact that both live and HK XZ17 alleviated virus-induced pulmonary fibrosis in HCT mice implies the structural components of the bacteria, rather than secreted soluble factors such as metabolites, were sufficient for effects on fibrosis. Since HK *L. johnsonii* offers the benefit of mitigating pulmonary fibrosis without the risk of causing infection in immunocompromised HCT recipients, subsequent studies focus on deciphering the mechanisms underlying the effects of the HK *L. johnsonii* XZ17 strain, henceforth referred to as HK *Lj* administration.

**Figure 2.**
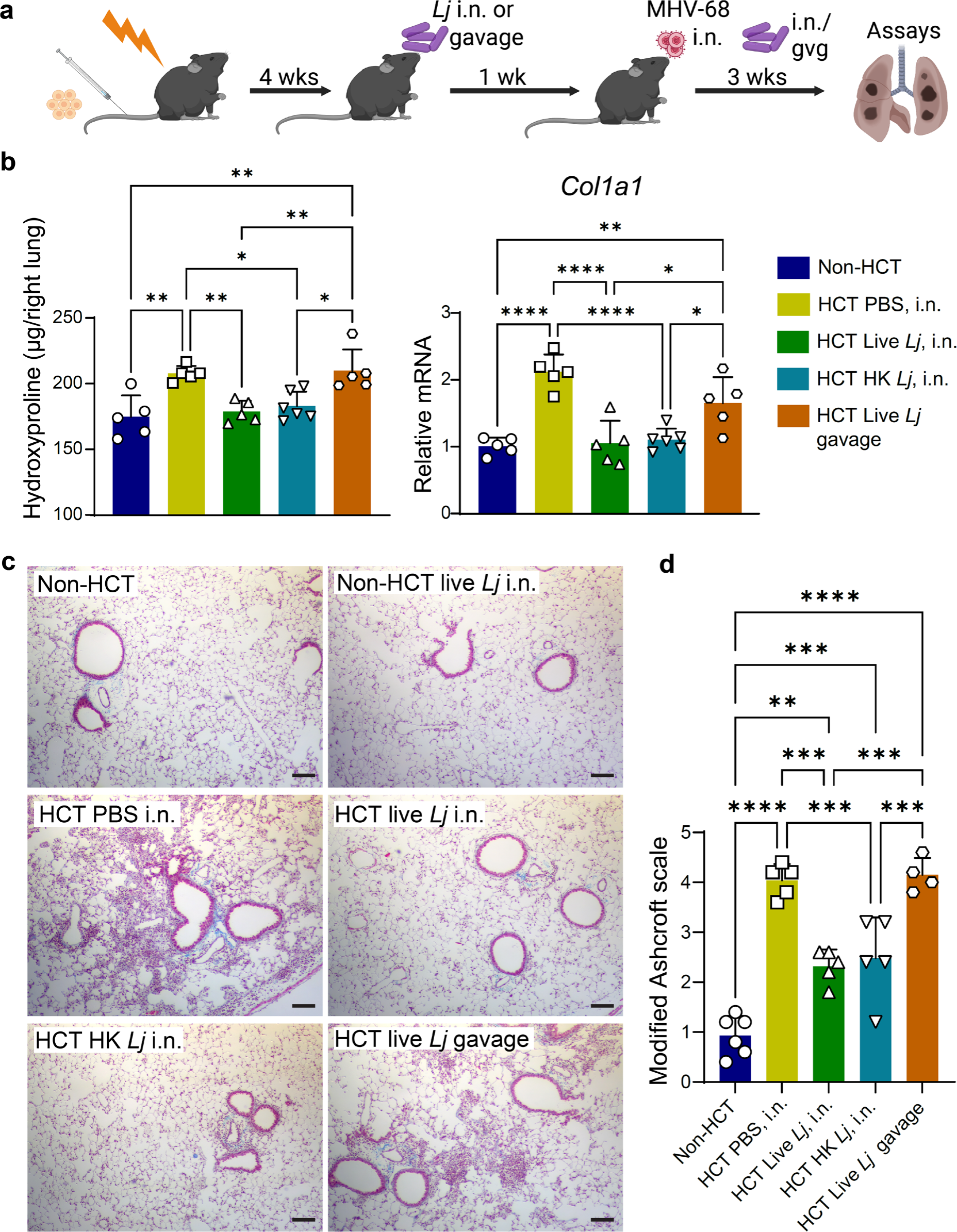
Intranasal administration of live or heat-killed *L. johnsonii* XZ17 reduces pulmonary fibrosis in HCT mice infected with MHV-68. a. Schematic of the experimental design (created with BioRender.com). C57BL/6J mice (n=5 to 6 per group) were lethally irradiated before transplantation with 5×10^6^ bone marrow cells from donor C57BL/6J mice. At 4 weeks post-transplantation, either 5×10^5^ CFU of live *L. johnsonii* XZ17 (Live *Lj*) or an equivalent amount of heat-killed *L. johnsonii* XZ17 (HK *Lj*) was administered intranasally to recipient mice at intervals of 2 to 3 days. Alternatively, mice were given 4×10^7^ CFU of live XZ17 via oral gavage at the same schedule. One week after the initial intranasal or oral administration of *L. johnsonii* XZ17, mice were infected intranasally with 5×10^4^ PFU of MHV-68, followed with continuous administration of XZ17, and lung tissues were collected at 21 days post-infection. **b**. Collagen content in the right lung was assessed by the hydroxyproline assay (left), and relative mRNA expression levels of collagen type I alpha 1 chain (*Col1a1*) in the left lung tissue was measured by qPCR (right). All samples were from MHV-68 infected mice. **c**. Representative images of lung tissue sections of MHV-68 infected mice stained with Masson’s Trichrome (scale bar, 100 μm). Collagen is stained blue. **d**. Modified Ashcroft scoring for fibrosis evaluation. Data are presented as mean ± SEM. Statistical significance is indicated by \**P*<0.05; \*\**P* < 0.01; \*\*\**P*<0.001; \*\*\*\**P* < 0.0001, as determined by one-way ANOVA with Tukey’s multiple-comparisons test. Results are representative of three independent experiments.

### Intranasal administration of HK *L. johnsonii* XZ17 exerts anti-inflammatory effects on macrophages and Th17 cells

Pulmonary fibrosis often develops following an inflammatory response to initial lung injury^30^. To determine whether intranasal administration of HK *Lj* mitigates pulmonary fibrosis by inhibiting inflammatory responses in the lungs of HCT mice infected with MHV-68, we conducted flow cytometry analyses on single-cell suspensions of lung immune cells at 7 dpi. The gating strategy utilized is presented in **Supplementary Fig. 2**. At 7 dpi, HK *Lj* administration did not significantly alter the relative or absolute numbers of many immune cell types, including T cells, neutrophils, natural killer cells (NK), resident and inflammatory monocytes (rMono and iMono), interstitial macrophages (IM), type II conventional dendritic cells (cDC2), monocyte-derived dendritic cells (MoDC), and eosinophils (Eos; **Supplementary Fig. 3a and b**). Tissue-resident alveolar macrophages (TRAM) were diminished in the lungs following HCT and infection, and administration with HK *Lj* did not restore their numbers. HK *Lj* administration led to a decrease in the numbers of B cells and type I conventional dendritic cells (cDC1) in the lungs of HCT mice at 7 dpi **(Supplementary Fig. 3a and b**). Intriguingly, HK *Lj* administration increased recruitment of monocyte-derived alveolar macrophages (MoAM) despite that HK *Lj* administration did not increase the expression of *Ccl2*, a chemokine that typically mediates the recruitment of monocytes and macrophages in the lung^31,32^ (**Supplementary Fig. 3c**). HK *Lj* administration did not significantly affect the replication of MHV-68 at 7dpi as evaluated by expression of the viral DNA polymerase gene (*pol*; **Supplementary Fig. 3c**). We observed a trend towards decreases in the mRNA levels of the inflammatory cytokine genes *Il6* and *Il1b* following HK *Lj* administration in the whole lung (**Supplementary Fig. 3c**). Considering that macrophages are key producers of inflammatory cytokines, and MoAMs are critically involved in the development of pulmonary fibrosis^32,33^, we enriched macrophages from the lung single-cell suspensions using microbeads conjugated to anti-F4/80 antibodies. We detected a significant reduction in the mRNA expression levels of the inflammatory cytokines *Il6*, *Il1b*, and *Il18* in macrophages from lungs of HCT mice that received HK *Lj*, along with significantly decreased *Tgfb1* mRNA levels (**Fig. 3a**). Of note, activated TGF-β1 is a significant contributing factor to the onset of pulmonary fibrosis^15,34,35^. In line with the concept that alternatively activated M2-type macrophages play an important role in promoting pulmonary fibrosis^7,36^, our findings indicate that HK *Lj* administration suppressed M2 phenotype, as evidenced by downregulation of *Arg1*, a marker for M2 macrophages, in the absence of an effect on expression of *Inos*, a marker for M1 macrophages (**Fig. 3a**). These findings indicate that, although HK *Lj* administration stimulates the recruitment of MoAMs into the lungs, HK *Lj* exerts anti-inflammatory effects by modulating macrophage phenotypes.

**Figure 3.**
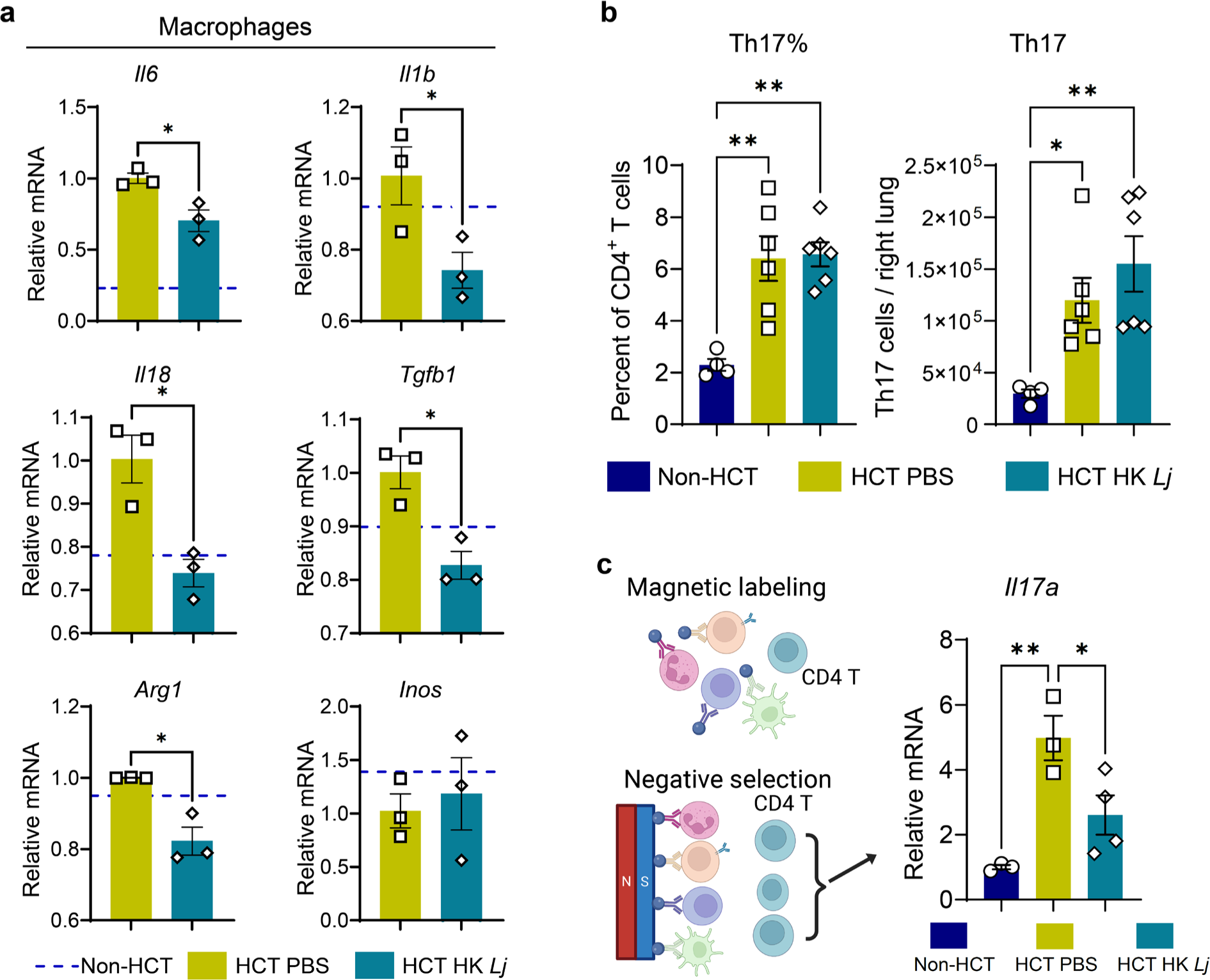
Intranasal administration of HK *L. johnsonii* XZ17 reduces inflammation in macrophages and Th17 cells at 7 days post infection with MHV-68. C57BL/6J mice (n=6 per group), with or without HCT, received intranasal administration of HK *Lj* or PBS every 2 to 3 days. All mice were then intranasally infected with MHV-68, and lung tissues were harvested at 7 dpi. **a**. Relative mRNA expression levels of a variety of cytokines in lung macrophages at 7 dpi with MHV-68. Macrophages were isolated from pooled single-cell suspensions of two lungs using microbeads conjugated with anti-F4/80 antibodies, and qPCR was used to assess relative mRNA expression levels. The dashed lines represent expression levels of each gene in lung macrophages from non-HCT mice infected with MHV-68. **b**. Flow cytometry was used to measure the proportion of Th17 cells (CD45^+^CD3^+^CD4^+^IL-17^+^) within the CD4^+^ T cell population (left, n=4 to 6) and absolute Th17 cell numbers (right) in the right lung. **c**. Schematic of the negative selection process used to enrich for untouched CD4^+^ T cells (left); qPCR analysis of *Il-17a* mRNA expression in isolated CD4^+^ T cells (n=3 to 4 mice, right). Data are presented as mean ± SEM. For **a**, statistical significance is denoted by *P<0.05, calculated using unpaired two-tailed Student’s *t* tests. For **b** and **c**, statistical significance is denoted by \**P*<0.05; \*\**P* < 0.01, determined by one-way ANOVA with Tukey’s multiple-comparisons test. Results are representative of two independent experiments.

**Supplementary Figure 2:**
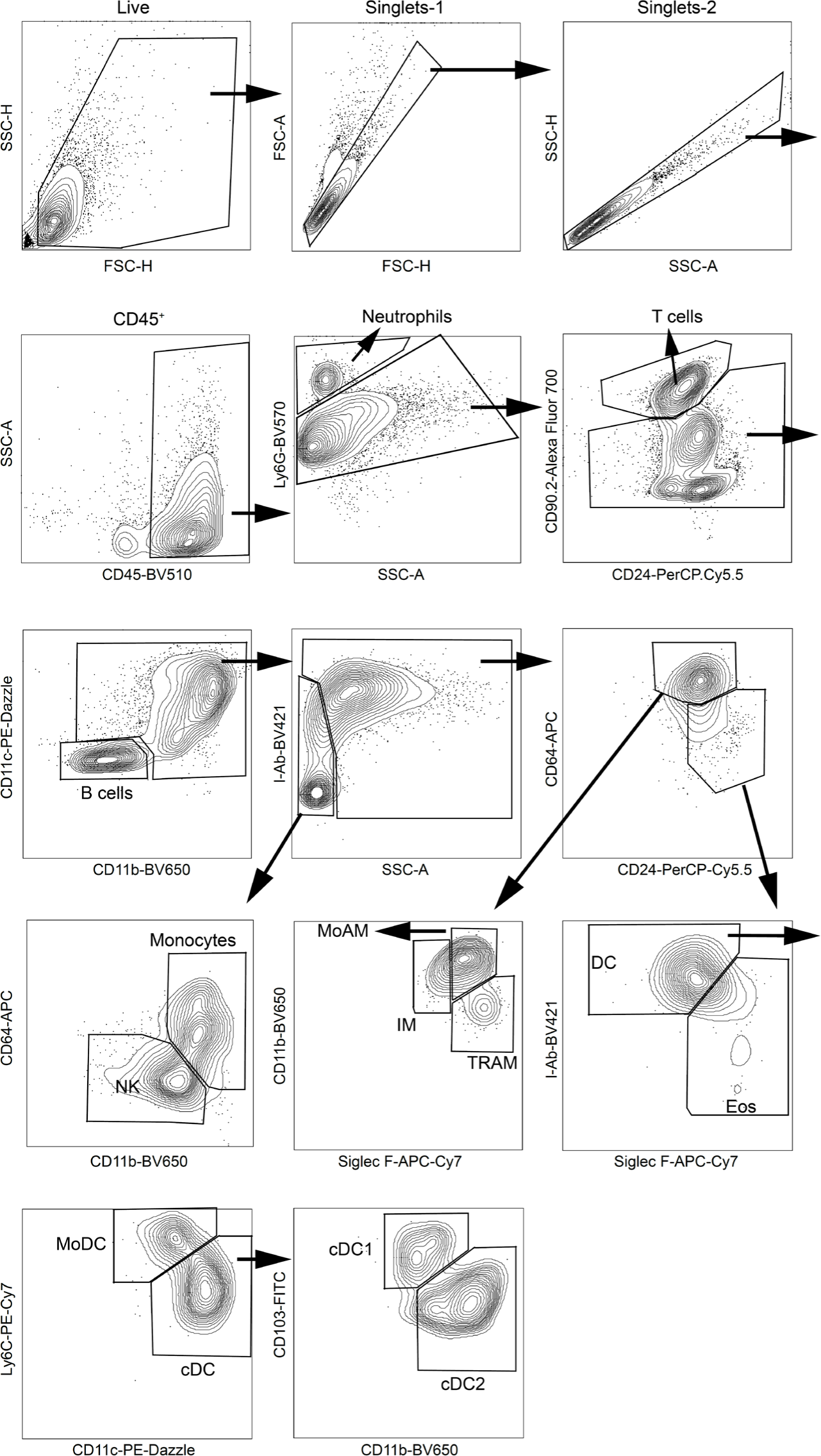
Gating strategy used for FACS analysis of populations of immune cells in the lungs at 7 days post infection with MHV-68.

**Supplementary Figure 3:**
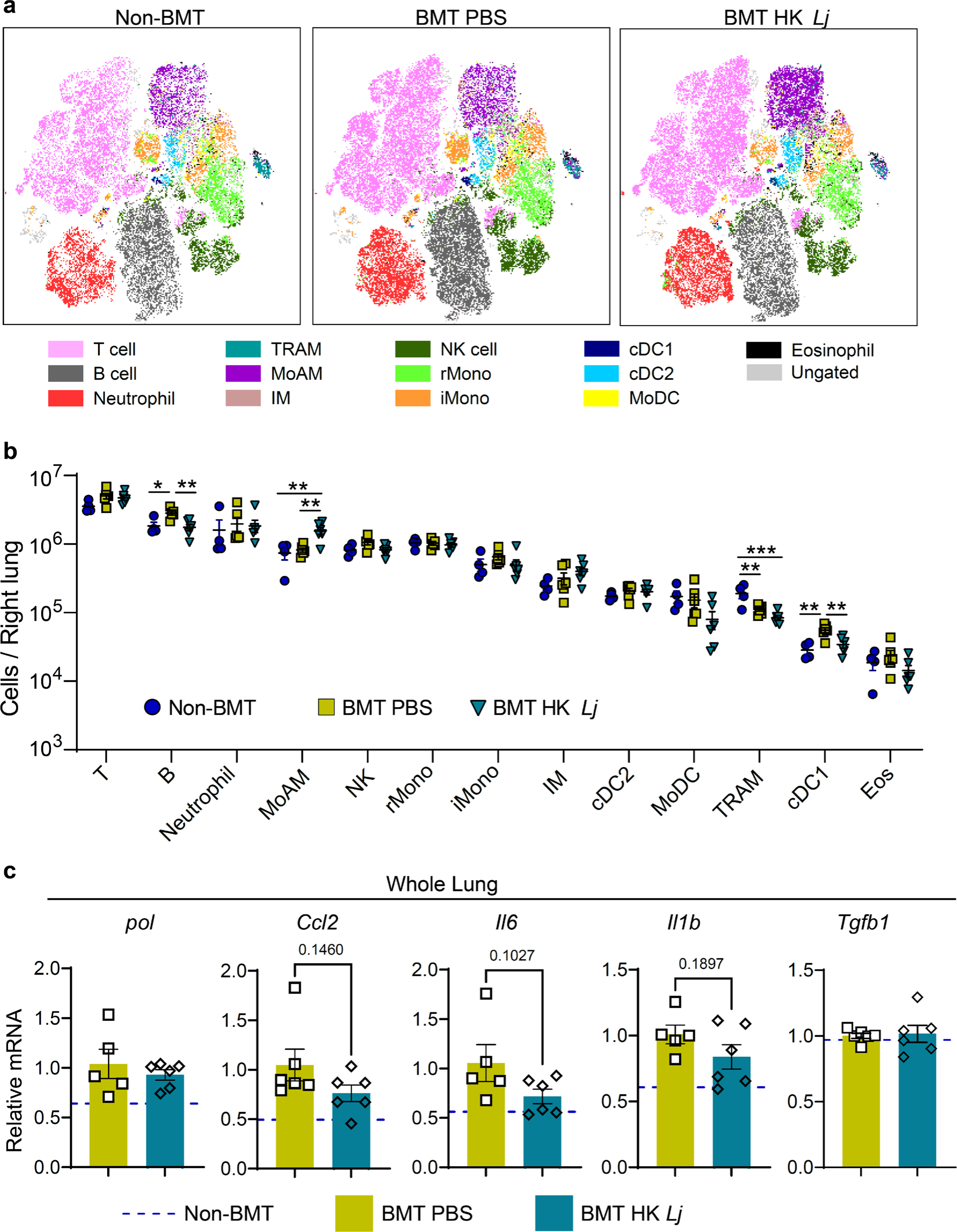
Flow cytometry analyses of lung immune cells and quantitative gene expression analysis in lung tissue following MHV-68 infection at 7 dpi. **a**. t-Stochastic Neighbor Embedding (t-SNE) visualization of lung immune cell populations based on flow cytometry markers. **b**. Quantification of absolute immune cell numbers for each identified lung immune cell type per right lung. **c**. Relative mRNA expression levels of MHV-68 viral polymerase (*pol*) and various cytokines in lung tissue assessed by qPCR. The dashed lines represent expression levels of each gene in lungs from non-HCT mice infected with MHV-68. Data are presented as mean ± SEM. For **b**, statistical significance is denoted by \**P*<0.05; \*\**P* < 0.01; \*\*\**P*<0.001, determined by one-way ANOVA with Tukey’s multiple-comparisons test. For **c**, *P*-values were calculated using unpaired two-tailed Student’s t-tests. Results shown are representative of two independent experiments.

We have established in previous research that lung Th17 cells (CD45^+^CD3^+^CD4^+^IL-17^+^) are increased in HCT mice after MHV-68 infection, and that IL-17A is essential for the development of pulmonary fibrosis in this viral-induced HCT fibrosis model^13,14,37^. However, to our surprise, the percentage and absolute numbers of Th17 cells in lungs at 7 dpi were not diminished by HK *Lj* administration, as determined by intracellular antibody staining and flow cytometry analysis (**Fig. 3b**). To investigate whether HK *Lj* administration could suppress Th17 cell expression of *Il17a* in HCT mice, we measured their *Il17a* mRNA expression in CD4^+^ T cells isolated from lungs (**Fig. 3c**). We found a significant decrease in *Il17a* expression in lung CD4^+^ T cells isolated from HK *Lj* -supplemented HCT mice at 7dpi (**Fig. 3c**). Thus, while HK *Lj* administration did not inhibit the polarization of Th17 cells in the lungs of HCT mice, it did reduce *Il17a* expression in those cells.

### HK *L. johnsonii* XZ17 administration enhances PD-L1 expression on the cell surfaces of cDC2s, MoDCs, and alveolar epithelial cells

The observed decrease in Th17 cell expression of *Il17a* in HCT mice that received HK *Lj* administration does not appear to be attributable to increased percentages of regulatory T cells (Treg), since we did not observe an effect of HK *Lj* administration on the proportion of RORγt^+^ Tregs or conventional Tregs in HCT mice (**Supplementary Fig. 4a and b**). This contrasts with a recent report in which an enhancement of immunosuppressive RORγt^+^ Tregs in the lungs was seen in naive mice after intranasal administration of a *Ligilactobacillus murinus* strain^38^.

We reasoned that HK *Lj* administration may enhance the expression of programmed death-ligand 1 (PD-L1) on the surface of antigen-presenting cells (APCs), which could in turn suppress Th17 cell activity via the PD-L1/programmed cell death-1 (PD-1) signaling pathway^39–41^. PD-L1 signaling from MHC II^+^ APCs can reduce cytokine production by CD4^+^ T cells when interacting through PD-1 receptors^42,43^. Indeed, we found that HK *Lj* administration increased expression of PD-L1 on the surface of type II conventional dendritic cells (cDC2s), MoDCs, and MHC II+ EpCAM+ alveolar epithelial cells (**Fig. 4a** and **b**). Among them, cDC2s exhibited the highest PD-L1 expression, while MoDCs and epithelial cells displayed relatively lower levels. Cell surface PD-L1 levels on TRAMs were increased following HCT and infection, but no additional increase was triggered by HK *Lj* administration at 7 dpi (**Fig. 4a** and **b**). Given that cDC2s play a critical role in directing the polarization of Th17 cells^14,44^, our findings imply that HK *Lj* administration may specifically target and inhibit Th17 cell activities. Despite the observed increases of PD-L1 expression on the antigen-presenting cells mentioned above, no decrease of CD4^+^ or CD8^+^ T cells were observed in HK *Lj*-supplemented HCT mice (**Supplementary Fig. 5a and 5b**). Notably, the percentage of PD-1^+^ cells in the CD4^+^ T cell and Th17 cell compartments at 7 dpi was increased in the lungs of HCT mice supplemented with HK *Lj* (**Fig. 4c**). We detected a similar increase in percentage of PD-1^+^ CD8^+^ T cells (**Supplementary Fig. 5c**). Thus, HK *Lj* administration increased the expression of PD-L1 on cDC2s, which potentially suppresses the expression of *Il17a* in Th17 cells in HCT lungs by augmenting PD-L1/PD-1 signaling.

**Figure 4.**
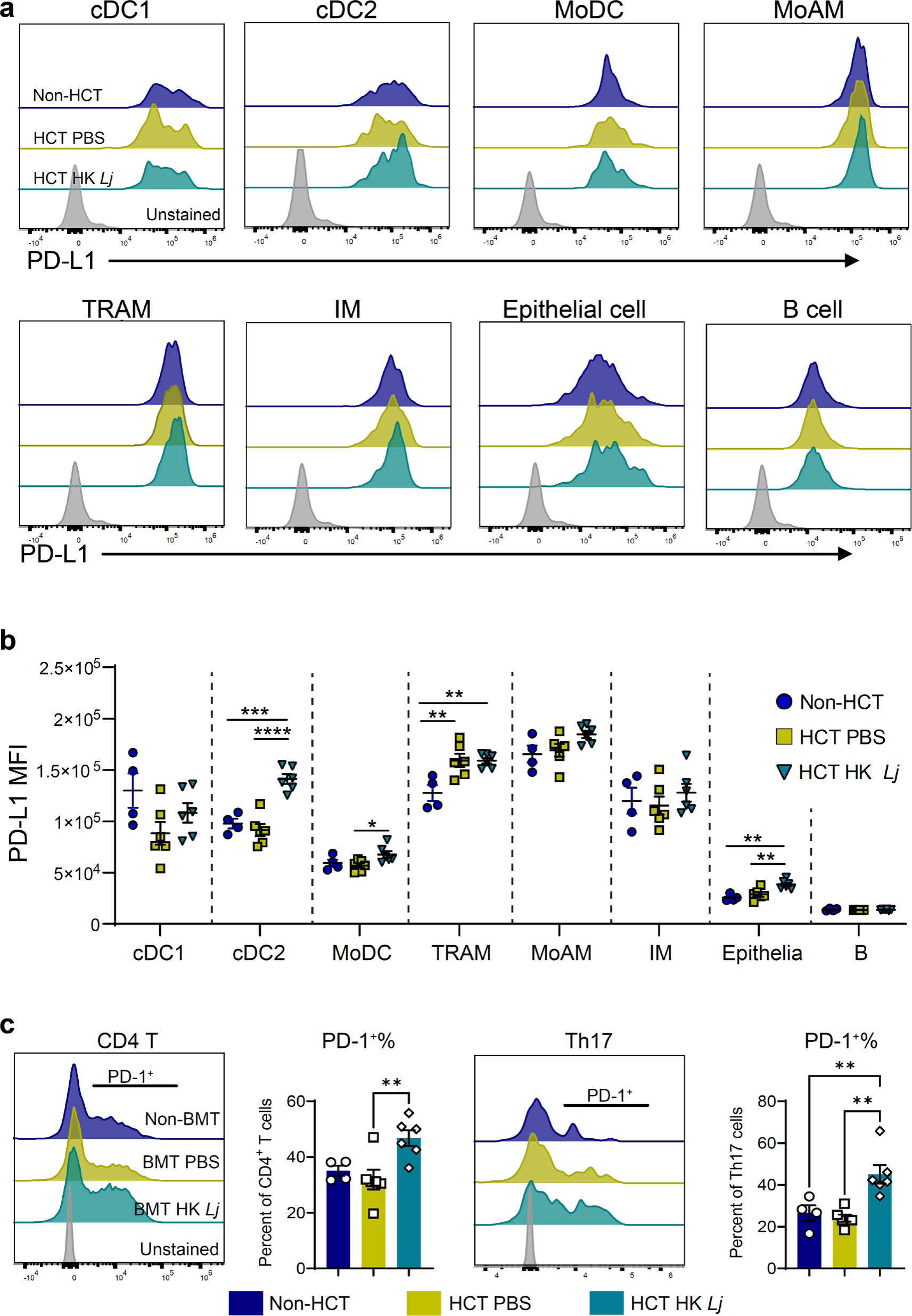
Intranasal administration of HK *L. johnsonii* XZ17 augments PD-L1 expression on the cell surfaces on cDC2s, MoDCs, and alveolar epithelial cells. C57BL/6J mice (n=6 per group), with or without HCT, received intranasal administration of HK *Lj* or PBS every 2 to 3 days. All mice were then intranasally infected with MHV-68, and lung tissues were harvested at 7 days post-infection for flow cytometry analysis. **a**. Representative flow cytometry histograms illustrating PD-L1 expression on MHC II^+^ cells, including dendritic cells, macrophages, epithelial cells, and B cells, at 7 days post-infection with MHV-68 (n=4 to 6). **b**. Mean fluorescence intensity (MFI) quantification of PD-L1 expression on dendritic cells, macrophages, epithelial cells, and B cells at 7 dpi (n=4 to 6). **c**. Left, representative histograms displaying PD-1 expression on CD4^+^ T cells accompanied by percentages of PD-1^+^ CD4^+^ T cells (n=4 to 6); Right, representative histograms presenting PD-1 expression on Th17 cells accompanied by percentages of PD-1^+^ Th17 cells (n=4 to 6). For **b** and **c**, data are presented as mean ± SEM, and statistical significance is represented by \*\**P* < 0.01; \*\*\**P*<0.001; \*\*\*\**P* < 0.0001, as determined by one-way ANOVA with Tukey’s multiple-comparisons test. Results are representative of two independent experiments.

**Supplementary Figure 4:**
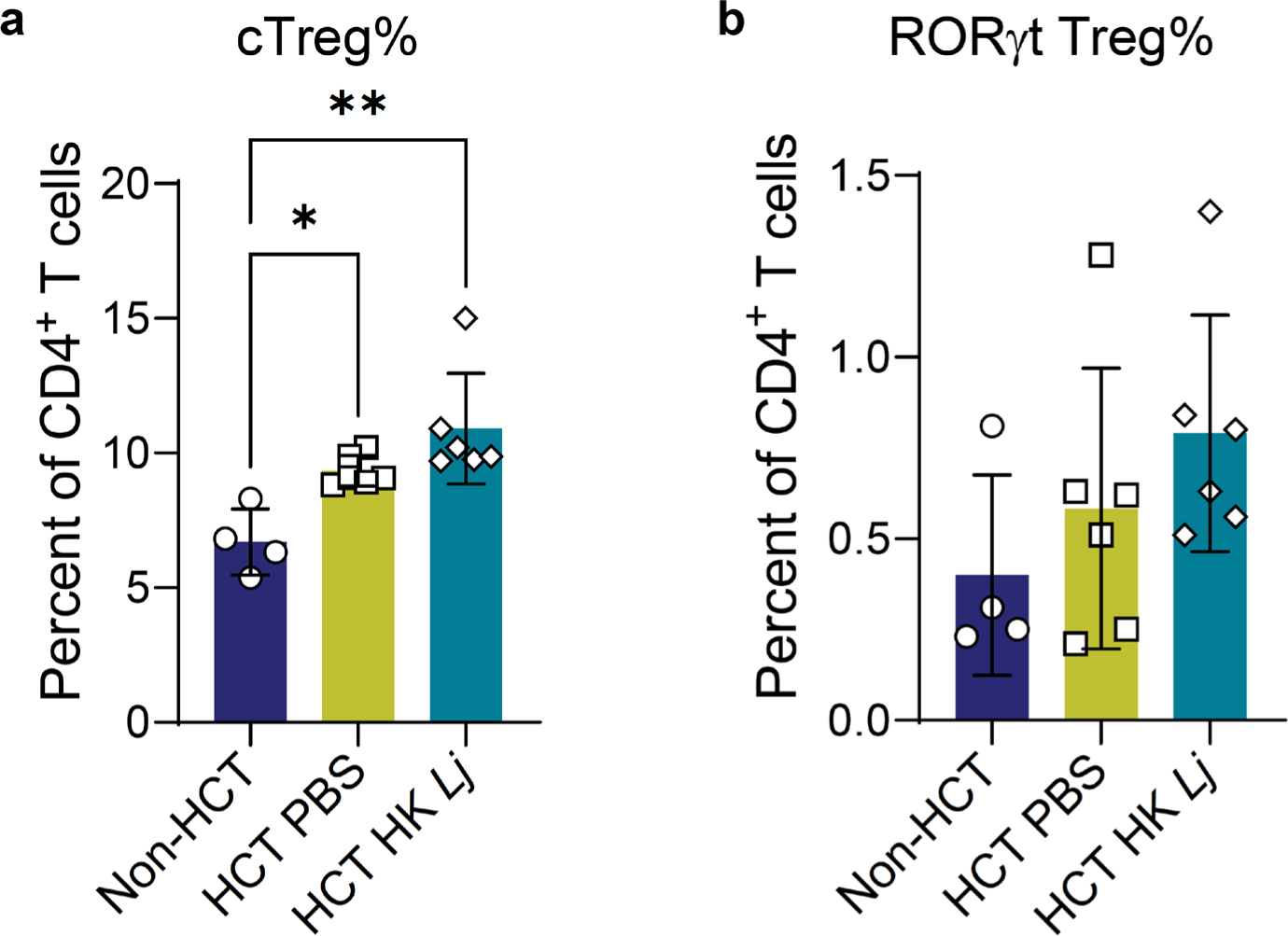
HK *L. johnsonii* XZ17 administration does not alter the frequency of regulatory T cells (Treg). **a**. Percentage of conventional Tregs (FoxP3^+^) within the CD4^+^ T cell population (n=4 to 6). **b**. Proportion of RORγt^+^ Tregs within the CD4^+^ T cell population (n=4 to 6), both evaluated by flow cytometry. Data are expressed as mean ± SEM. Statistical significance is indicated by \**P*<0.05; \*\**P* < 0.01, as determined by one-way ANOVA with Tukey’s multiple-comparisons test. The results are representative of two separate experiments.

### HK *L. johnsonii* XZ17 enhances the expression of PD-L1 on bone marrow-derived dendritic cells, which subsequently inhibits iTh17 cell production of IL-17A

The results from the *in vivo* studies described above indicate that HK *Lj* administration leads to the upregulation of PD-L1 on the surface of cDC2s in HCT mice. To investigate if HK *Lj* directly enhances dendritic cell surface expression of PD-L1, we generated bone marrow-derived dendritic cells (BMDCs) from a C57BL/6J male mouse (wild type, WT). Granulocyte-macrophage colony-stimulating factor (GM-CSF) was used to induce the differentiation of bone marrow progenitor cells into dendritic cells with phenotypic similarities to cDC2s and MoDCs^45^. These BMDCs were cultured with HK *Lj* at a 1:2 ratio for 24 hours (**Fig. 5a**). A significant increase in PD-L1 expression on BMDCs was observed following exposure to HK *Lj* (**Fig. 5b** and **c**). To evaluate whether the upregulation of PD-L1 on BMDC represses Th17 cell secretion of IL-17A, we co-cultured untreated or HK *Lj*-treated BMDCs with induced Th17 (iTh17) cells, which were differentiated from naive CD4^+^ T cells from the spleens of WT or PD-1 knockout (PD-1KO) mice^46^ (**Fig. 5d**). For iTh17 cells derived from WT mice, a notable reduction in IL-17A secretion occurred when co-cultured with HK *Lj*-treated BMDCs, compared to iTh17 cells co-cultured with untreated BMDCs. Conversely, iTh17 cells derived from PD-1KO mice did not exhibit suppressed IL-17A production following exposure to HK *Lj*-treated BMDCs. Instead, IL-17A production was significantly increased when PD-1KO iTh17 cells were co-cultured with HK *Lj*-treated BMDCs (**Fig. 5e**). These results strongly suggest that HK *Lj* exposure suppresses iTh17 cell production of IL-17A through engagement of the PD-L1/PD-1 signaling axis.

**Figure 5.**
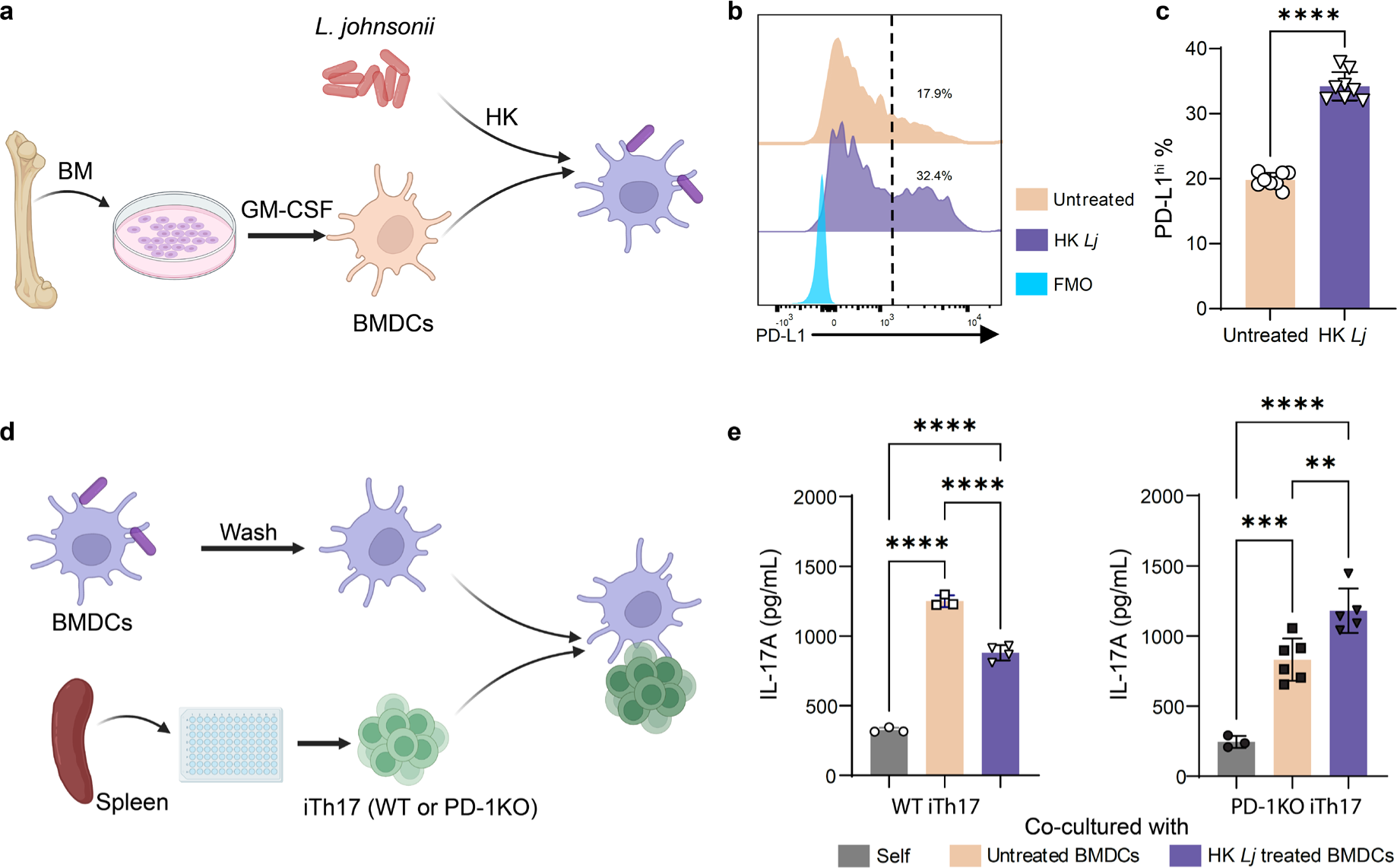
HK *L. johnsonii* XZ17 enhances PD-L1 expression on BMDCs and suppresses IL-17A production in iTh17 cells via PD-L1/PD-1 signaling. a. A diagram depicting the procedure to derive BMDCs from bone marrow cells and their subsequent co-culture with HK *Lj* (created with BioRender.com). **b**. Flow cytometry histograms showing PD-L1 expression on BMDCs. Cells to the right of the dashed line are high expressers of PD-L1 (PD-L1^hi^), with numbers indicating the percentage of these cells. **c**. Proportion of PD-L1^hi^ BMDCs based on HK *Lj* treatment (n=8 per group). **d**. Diagram illustrating the generation of iTh17 cells and their co-culture with HK *Lj*-treated BMDCs (created with BioRender.com). **e**. ELISA was used to measure IL-17A concentrations in co-culture supernatants. The left panel shows co-cultures with BMDCs and WT iTh17 cells; the right panel shows co-cultures with BMDCs and PD-1KO iTh17 cells. N = 3 to 6 culture supernatants. Data are expressed as mean ± SEM. For **c**, statistical significance is denoted by \*\*\*\**P* < 0.0001, determined using unpaired two-tailed Student’s *t* tests. For **e**, statistical significance is indicated by \*\**P* < 0.01; \*\*\**P*<0.001; \*\*\*\**P* < 0.0001, using one-way ANOVA with Tukey’s multiple-comparisons test. The depicted results are representative of two independent experiments.

**Supplementary Figure 5:**
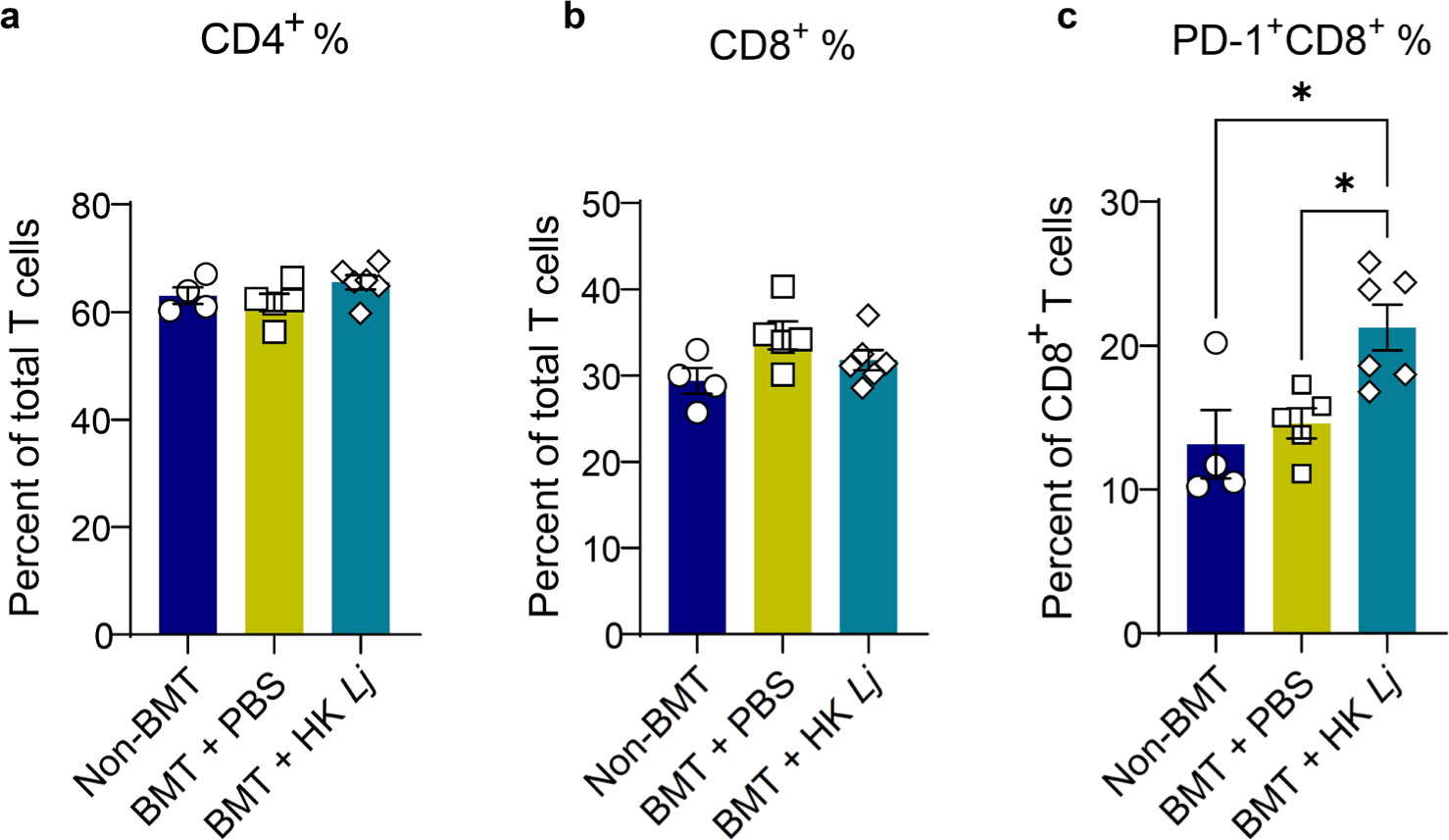
HK *L. johnsonii* XZ17 administration does not alter the frequency of CD4 and CD8 T cells. **a**. Percent of CD4+ T cells. **b**. Percent of CD8+ T cells. **c**. Percent of PD-1+ CD8+ T cells. Data are expressed as mean ± SEM. Statistical significance is indicated by \**P*<0.05, as determined by one-way ANOVA with Tukey’s multiple-comparisons test. The results are representative of two separate experiments.

### HK *L. johnsonii* XZ17 alleviates pulmonary fibrosis in MHV-68-infected HCT mice in a PD-L1/PD-1 signaling-dependent manner

Taking into account the increase in PD-L1 expression on antigen presenting cells following HK *Lj* treatment in HCT mice, as well as the *in vitro* evidence indicating that HK *Lj* utilizes the PD-L1/PD-1 signaling axis to reduce IL-17A secretion from iTh17 cells, we hypothesized that the alleviation of pulmonary fibrosis in MHV-68-infected HCT mice following intranasal delivery of HK *Lj* is mediated by the activation of the PD-L1/PD-1 signaling pathway. To test this hypothesis, we developed PD-1KO HCT mice, wherein bone marrow cells from PD-1KO mice were transplanted into irradiated syngeneic recipients. In these PD-1KO mice, T cells are unresponsive to PD-L1 signaling from PD-L1-expressing cells. WT HCT mice began to experience weight loss after 7 dpi with MHV-68, and intranasal administration of HK *Lj* in WT HCT mice trended toward mitigating this MHV-68-induced weight loss, with effects becoming noticeable starting at 11 dpi (**Fig. 6a**). Compared to WT HCT mice, PD-1KO HCT mice experienced more pronounced weight loss starting at 7 dpi with MHV-68 and succumbed to the infection by 12 dpi (**Fig. 6a** and **b**). These outcomes are consistent with the protective role of PD-1 in preventing lethal immunopathology during acute viral infections^47^. HK *Lj* administration attenuated both weight loss and mortality in the infected PD-1KO HCT mice (**Fig. 6a** and **b**), suggesting that some protective effects of HK *Lj* are PD-1-independent. However, while administration with HK *Lj* improved weight loss and survival in infected PD-1KO HCT mice, it failed to mitigate lung fibrosis (as detected by hydroxyproline assay) to the same extent as was observed in WT HCT counterparts (**Fig. 6c**). It should be noted that the seemingly low levels of lung collagen content in infected PD-1KO HCT mice without HK *Lj* administration might be attributed to evaluation at an earlier endpoint (11-12 dpi) in these mice, compared to the other groups assessed at 15 dpi (**Fig. 6c**). These results suggest that the PD-1 pathway plays an essential role in mediating the anti-fibrotic effect of HK *Lj* administration.

**Figure 6.**
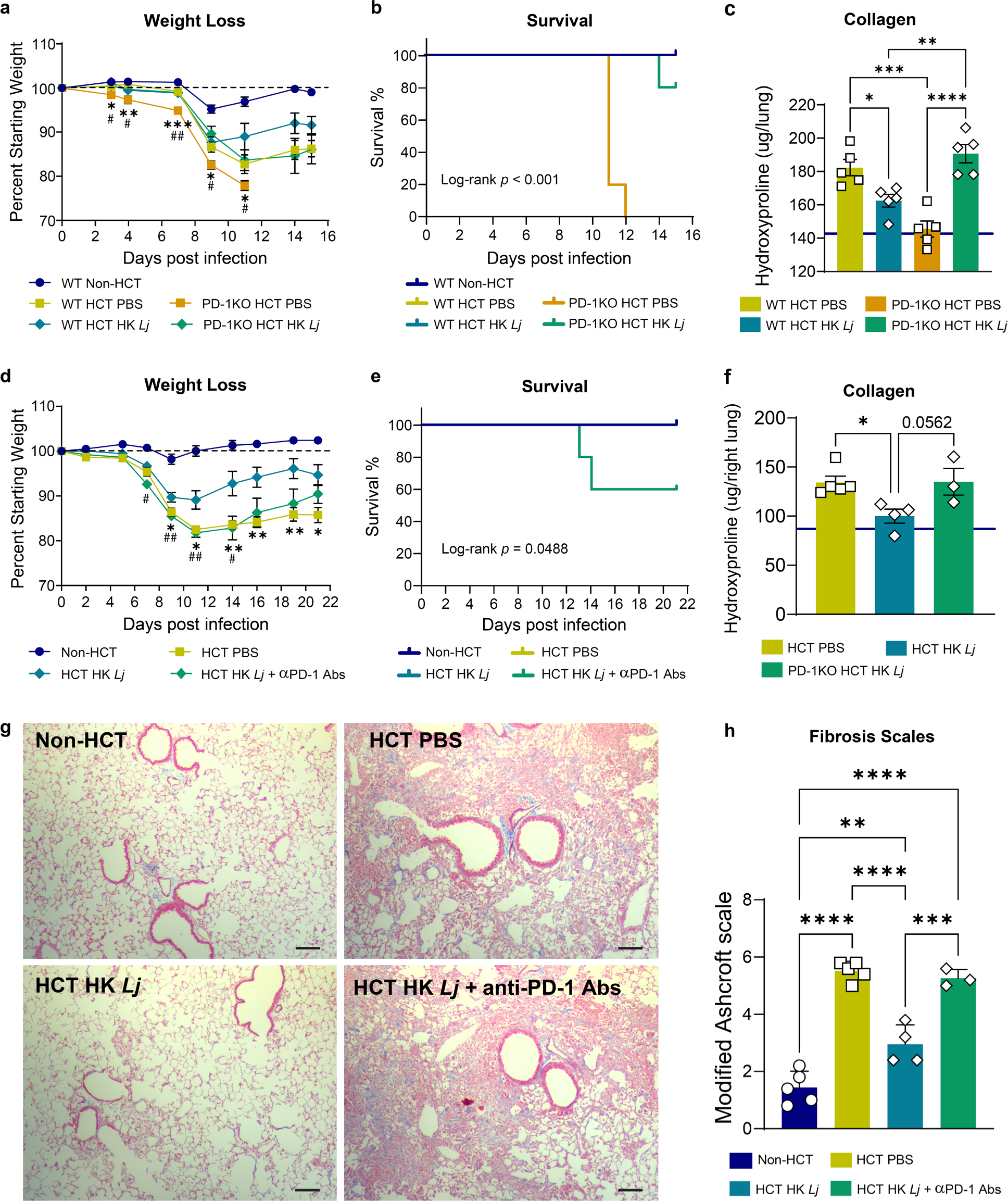
The PD-L1/PD-1 signaling axis is required for HK *L. johnsonii* XZ17 to alleviate pulmonary fibrosis in HCT mice after MHV-68 infection. **a-c**. Effects of HK *Lj* administration on HCT mice reconstituted with PD-1KO bone marrow cells. C57BL/6J mice (n=5 per group) underwent bone marrow transplantation with cells from either C57BL/6J mice (WT HCT) or PD-1KO mice (PD-1KO HCT). Beginning at 4 weeks post-transplantation, each mouse was administered HK *Lj* or PBS intranasally at 2-to 3-day intervals. Subsequently, all mice were infected intranasally with MHV-68. Results are representative of three independent experiments. **a**. Weight loss in mice after MHV-68 infection. **b**. Survival curves. **c**. Hydroxyproline assay measurements of collagen content in whole lungs. Lungs were harvested at either 15 dpi or upon reaching the moribund stage. **d-h**. Effects of HK *Lj* administration on PD-1 neutralized HCT mice. WT HCT mice were administrated with 100 μg rat anti-mouse PD-1 neutralizing antibodies (29F.1A12) or PBS (n=3 to 5 per group). **d**. Weight loss in mice after MHV-68 infection. **e**. Survival curves. **f**. Collagen quantification in right lungs using the hydroxyproline assay. **g**. Masson’s Trichrome staining of lung tissue sections for visual assessment of fibrosis (scale bar, 100 μm). Collagen is stained blue. **h**. Evaluation of fibrosis using the modified Ashcroft scale. Data are presented as mean ± SEM. For panel **a**, significant differences in body weight changes: WT HCT PBS versus PD-1KO HCT PBS, \**P*<0.05; \*\**P* < 0.01; \*\*\**P* < 0.001; PD-1KO HCT PBS versus PD-1KO HCT HK *Lj*, ^#^*P*<0.05; ^##^*P* < 0.01, using unpaired two-tailed Student’s t-tests. For panel **d**, significant differences in body weight changes: HCT PBS versus HCT HK *Lj*; \**P*<0.05; \*\**P* < 0.01, ^#^*P*<0.05; HCT PBS versus HCT HK *Lj +* αPD-1 Abs, ^##^*P* < 0.01, using unpaired two-tailed Student’s t-tests. For panels **c**, **f**, and **h**, statistical significance is denoted by \**P*<0.05; \*\**P* < 0.01; \*\*\**P*<0.001; \*\*\*\**P* < 0.0001, as determined by one-way ANOVA with Tukey’s multiple-comparisons test.

We extended our investigation into the role of PD-L1/PD-1 signaling in pulmonary fibrosis using an anti-PD-1 neutralizing antibody that has been previously shown to be effective in promoting antitumor immunity^48–50^. Intranasal administration of HK *Lj* ameliorated MHV-68-induced weight loss in HCT mice; however, HK *Lj* treatment did not significantly impact weight loss in PD-1 neutralized HCT mice post-MHV-68 infection (**Fig. 6d**). Instead, PD-1 neutralization was associated with increased mortality (**Fig. 6e**). Consistent with our findings in PD-1KO HCT mice, PD-1 neutralization in WT HCT mice inhibited effects of HK *Lj* administration on pulmonary fibrosis assessed by measuring lung collagen content (**Fig. 6f**) and evaluating histological evidence of fibrosis (**Fig. 6g** and **h**). Collectively, we have demonstrated that the PD-L1/PD-1 signaling axis mediates the effect of HK *Lj* in reducing pulmonary fibrosis in HCT mice following viral infection.

## Discussion

We have demonstrated that intranasal administration of low-dose nonviable *L. johnsonii* XZ17, a bacterium often found in normal mouse lungs but diminished following HCT, alleviated pneumonitis and pulmonary fibrosis in HCT mice infected with MHV-68. This effect was mediated by the upregulation of PD-L1 on cDC2s, which subsequently dampened IL-17A production in Th17 cells. Furthermore, although administration of HK *Lj* increased the recruitment of MoAMs, it reduced pro-inflammatory cytokine production by lung macrophages. Interestingly, both live and inactivated *L. johnsonii* can alleviate virus-induced pulmonary fibrosis in HCT mice.

Nonviable microbial entities, with or without their metabolites, are recognized as postbiotics by the International Scientific Association for Probiotics and Prebiotics^51^. While live lactobacilli are typically safe when used as probiotics in healthy individuals, they may pose an opportunistic infection risk to immunocompromised hosts, such as HCT recipients^52–54^. Moreover, many strains of *L. johnsonii* harbor tetracycline resistance genes, which pose a risk of horizontal gene transfer of antibiotic resistance^55^. A recent clinical trial demonstrated that oral administration of heat-killed *L. johnsonii* No. 1088 was safe and helped improve temporary heartburn symptoms in healthy individuals^56^. In preclinical models, non-viable *L. johnsonii* has been shown to inhibit the growth of *Helicobacter pylori*^57^, protect against diet-induced obesity^58^, and suppress IL-12 production by macrophages^59^. The modulatory effects of nonviable lactobacilli on host immune responses are attributed to distinct non-soluble bacterial components such as peptidoglycan, teichoic acids, cell wall polysaccharides, and surface proteins^60^. Because nonviable bacteria such as HK *L. johnsonii* are likely to be safe in immunocompromised hosts, the effects of HK *L. johnsonii* that we defined in this study could serve as the basis for novel approaches to manage pulmonary complications following HCT.

Our study indicates that the soluble components of *L. johnsonii*, may exert only a minimal impact in the HCT mouse model. However, other research has demonstrated that the soluble components of *L. johnsonii*, such as metabolites and extracellular vesicles, possess distinct bioactivities within their hosts. For example, oral administration of live, but not heat-killed, *L. johnsonii* MR1 increases levels of serum docosahexaenoic acid, affecting immune responses during RSV infections in the lungs^27^. Extracellular vesicles from *L. johnsonii* N6.2 have been proposed to be critical in reducing the onset of Type 1 diabetes in Biobreeding Diabetes-Prone (BBDP) rats^61–63^. These observations highlight the various ways in which different components of *L. johnsonii* could provoke diverse biological responses.

*L. johnsonii* is commonly found in the lungs of healthy non-HCT mice, but these bacteria are likely to be transient visitors rather than permanent residents. We detected approximately a two-log reduction in lung culturable bacterial counts just one day after inoculating HCT mice with live *L. johnsonii* XZ17. This rapid reduction likely results from the challenging environment in the lungs, with factors such as limited nutrient availability, high oxygen stress, and the relentless clearance by alveolar macrophages. The lung microbiome is often a dynamic, low biomass community with similar community structures to that of the upper respiratory tract^64,65^. Bacteria are primarily introduced into the lungs through subclinical microaspiration along a mouth-lung migration route, rather than dispersion along contiguous respiratory mucosa^65,66^. Mice exhibit coprophagic behavior, which could promote the transfer of microbiota between the gut and the lungs. In line with this, our prior research revealed a concurrent reduction in the relative abundances of *Lactobacillus* genus members in both the lung and gut microbiomes of HCT mice^7^. These findings imply that a significant gut-lung microbiome interaction might be present during the development of pulmonary fibrosis.

However, oral gavage of *L. johnsonii* XZ17 in HCT mice did not significantly reduce the severity of pulmonary fibrosis, suggesting that XZ17 may have a more potent local effect in the lungs, while its influence through the gut-lung axis following oral administration may be negligible or too subtle to detect. The lack of effectiveness of oral administration in our HCT mouse model could be context-specific, possibly due to radiation-induced gut injury and subsequent dysbiosis post-HCT that might hinder lactobacilli colonization in the gut^7^. This stands in contrast to our recent findings on bleomycin-induced pulmonary fibrosis where oral antibiotics provided profound protection^67^, implying that the gut microbiome has a significant role in the development of bleomycin-induced pulmonary fibrosis.

Interstitial lung diseases (ILDs), which are characterized by inflammation and fibrosis in the lung interstitium, encompass approximately 200 distinct conditions^68^, implying a diverse etiology and pathogenesis. In IPF, the most severe and prevalent form of pulmonary fibrosis, studies have identified elevated levels of PD-L1 in serum and on the surface of epithelial cells, as well as increased PD-1 expression on T cells, although those studies were limited by their retrospective nature and small cohort sizes^69,70^. Correspondingly, in animal models of IPF, such as those induced by bleomycin, augmented PD-L1 and PD-1 expression levels have been reported, suggesting a role for the PD-L1/PD-1 pathway in the development of IPF-like fibrosis^69,70^. Those observations stand in contrast to the findings of our study, in which stimulation of PD-L1/PD-1 signaling by HK *L. johnsonii* XZ17 attenuated pulmonary fibrosis. This difference suggests the presence of distinct contributors to the pathogenesis of bleomycin-induced and herpesvirus-induced pulmonary fibrosis. Supporting this distinction, CCL2/CCR2 signaling, which is crucial for monocyte recruitment to the lungs, is required for the development of bleomycin-induced pulmonary fibrosis^32^ but helps control the severity of herpesvirus-induced pulmonary fibrosis in HCT mice^71^.

We observed that *L. johnsonii* promoted the expression of PD-L1 on dendritic cells, which in turn suppressed T cell functions. This finding prompts the need for caution when considering immune checkpoint blockade (ICB) therapies that target PD-L1/PD-1 signaling. A retrospective analysis of a large cohort of melanoma patients revealed that high dietary fiber intake correlated positively with ICB efficacy, but the consumption of probiotics abolished the benefits conferred by dietary fiber^72^. Moreover, mice that received probiotics containing *L. rhamnosus* GG or *Bifidobacterium longum* exhibit a diminished antitumor response to anti-PD-L1 therapy, leading to larger tumors with fewer infiltrated IFN-γ^+^ CD8^+^ T cells and Th1 cells^72^. Other studies have reported mixed effects of probiotics on ICB treatments across preclinical and clinical cancer settings^73–76^. Similarly, *L. rhamnosus* GG administration did not significantly affect the development of acute graft-versus-host disease (aGVHD) in a small cohort of allo-HCT patients^77^. However, a more recent study suggested that a regimen combining seven probiotic bacterial strains with a prebiotic significantly decreases both the incidence and severity of aGVHD, potentially via the induction of CD4^+^CD25^+^Foxp3^+^ regulatory T cells^78^. This discrepancy may relate to the involvement of formulations with different specific probiotic species. A recent study showed that oral administration with a live mouse isolate of *L. johnsonii* enhances the response to anti-PD-1 cancer therapy for various cancers by promoting the production of indole-3-propionic acid in the gut, which in turn activates progenitor exhausted CD8^+^ T cell^79^. These findings underline the necessity for more precise inquiries into the context-specific mechanisms of immune modulation by lactobacilli that contribute to improved health outcomes.

Overall, this study presents a promising and potentially safe therapeutic approach that involves the respiratory delivery of postbiotics to mitigate severe pulmonary complications in HCT recipients. It reveals an underlying mechanism by which heat-inactivated *L. johnsonii* suppresses the activities of Th17 cells through the promotion of the PD-L1/PD-1 signaling pathway from dendritic cells to helper T cells. This research paves the way for numerous future investigative opportunities, such as exploring the origins of the lung microbiome and developing targeted therapies based on the lung microbiome, identifying the active components of *L. johnsonii* that modulate dendritic cell responses, and elucidating the mechanisms by which dendritic cells interact with postbiotics.

## Methods

### Mice

C57BL/6J (B6) and B6.Cg-Pdcd1^tm1.15hr^/J (PD-1KO) mice were purchased from the Jackson Laboratory (Bar Harbor, ME). The mice were housed in a specific pathogen-free facility at University of Michigan. Upon reaching the predetermined endpoints, mice were euthanized utilizing CO2 (Cryogenic Gases, Detroit, MI), and organs were harvested for subsequent cell preparation or assay procedures. All procedures involving mice were approved by the University of Michigan’s Institutional Animal Care & Use Committee, as per the protocol number PRO00010437.

### Syngeneic hematopoietic cell transplantation and MHV-68 Infection

Recipient B6 mice (7∼9 weeks old) received a total body irradiation of 13 Gy by a ^137^Cs irradiator, administered in two split doses three hours apart. Subsequently, 5 x 10^6^ whole bone marrow cells, harvested from B6 or PD-1KO mice, were infused via tail vein injection. At 5 weeks post-HCT, the mice were inoculated with MHV-68 (VR-1465, ATCC). To do that, both non-HCT and HCT mice were anesthetized using a mixture of ketamine and xylazine, followed by the intranasal administration of 5 x 10^4^ PFU MHV-68.

### Isolation and identification of *L. johnsonii*

A lung from a healthy naïve adult B6 mouse was homogenized in 1 ml PBS under aseptic conditions. Lung homogenate was then streaked onto Lactobacillus-selective de Man, Rogosa, and Sharpe (MRS) agar (BD Biosciences) for 24 hours at 37°C under an aerobic environment supplemented with CO2. Pure cultures of eight colonies were established by re-streaking and subsequently maintained as frozen stocks. Bacterial identification was performed through PCR amplification of a segment of the 16S ribosomal RNA gene employing the degenerate primers D88 and E94^19^. Sequencing analysis confirmed that all isolates were genetically identical, displaying 100% sequence homology with the 16S rRNA gene sequence of *Lactobacillus johnsonii*^20^. A single isolate, designated *L. johnsonii* XZ17, was selected for full genomic sequencing (Ravi et al. accepted by *Microbial Resource Announcements*). The genome of XZ17 was deposited under GenBank accession number CP151183.1.

### Phylogenetic analysis on *L. johnsonii* genomes

Phylogenomic analysis was carried out on complete genome sequences of 17 *L. johnsonii* strains sourced from a variety of vertebrate hosts including humans, rodents, swine, and poultry. These strains are BS15, ATCC 33200, IDCC9203, NCC 533, CH32, FI9785, UMNLJ21, UMNLJ22, ZLJ010, PF01, DPC 6026, Byun-jo-01, MR1, MT4, PC38, and NCK2677. The type strain ATCC 33323 of *L. gasseri*, the species most closely related to *L. johnsonii*, was utilized as the outgroup external reference. Average nucleotide identity (ANI) analyses of whole genomes were conducted utilizing fastANI^80^. The resulting pairwise ANI scores were used to create a Euclidean distance matrix, from which a phylogenetic tree was constructed using hierarchical clustering in R. The final phylogenetic tree was visualized using Phylo.io^81^, enabling elucidation of the genetic relationships among the strains.

### Preparation and administration of *L. johnsonii*

For the preparation of *L. johnsonii* XZ17 stocks, 100 mL of MRS broth was inoculated with the bacterium from a frozen glycerol stock and cultured statically overnight at 37°C. The stationary-phase cultures were pelleted by centrifugation at 1,500 x g for 5 minutes at 4°C and subsequently resuspended in 50 mL of a 1:1 (vol/vol) solution of MRS broth and 50% glycerol. Aliquots of 500 μL of the bacterial suspension were stored at −80°C. The viable cell counts of the prepared frozen glycerol stocks were determined as 1 x 10^9^ CFU/vial through serial dilution.

To administer live *L. johnsonii* XZ17, stocks were rapidly thawed at 37°C, pelleted by by centrifugation at 1,500 x g for 2 minutes, and washed with PBS to eliminate excess glycerol. The washed cells were resuspended in sterile PBS to a final concentration of 1 x 10^4^ CFU/μL for intranasal delivery (50 μL/mouse), or to a concentration of 4 x 10^5^ CFU/μL for oral gavage (100 μL/mouse). Prior to intranasal administration, mice were sedated with isoflurane. The viability of *L. johnsonii* was confirmed by plating the remaining suspension on MRS agar throughout the experiment.

To generate heat-killed (HK) *L. johnsonii* XZ17 stocks, a vial of the frozen stock was quickly thawed at 37°C, washed with sterile PBS, resuspended in 1 mL of PBS, and then subjected to a 70°C water bath for 15 minutes to inactivate the bacteria. The heat-killed cells were pelleted, resuspended in sterile PBS to a final concentration equivalent to 1 x 10^6^ CFU/μL before heat treatment, and aliquoted into 15 μL vials, which were then stored at −80°C. The inactive status of the HK *L. johnsonii* stocks was confirmed by an overnight culture on MRS agar. For intranasal administration of HK *L. johnsonii*, 1.5 mL of PBS was added to a stock vial to prepare the inoculum, and 50 μL per mouse was administered intranasally.

### PD-1 blockade in HCT mice

Mice that underwent the HCT procedure and received HK *L. johnsonii* supplementation were intraperitoneally administered 100 µg of rat anti-PD-1 monoclonal neutralizing antibodies (clone 29F.1A12, BioXCell). The administration was carried out at intervals of 2 to 3 days, beginning at 2 days post-infection (dpi) with murine gammaherpesvirus 68 (MHV-68) and continuing through 21 dpi.

### RNA extraction and quantitative PCR analysis

RNA was isolated from lung tissues or specific cell types using TRIzol reagent (Invitrogen, Thermo Fisher Scientific) according to the instructions provided by the manufacturer. The concentration and purity of the extracted RNA were evaluated using a NanoDrop Lite Spectrophotometer (Thermo Fisher Scientific). Gene expression levels were quantified by reverse transcription quantitative PCR (RT-qPCR) using the Luna Universal Probe One-Step RT-qPCR Kit (New England Biolabs), and being conducted on a QuantStudio 3 Real-Time PCR System (Applied Biosystems). Detailed information on the primer and probe sets used in this study can be found in Supplemental Table 1.

**Supplemental Table 1.**
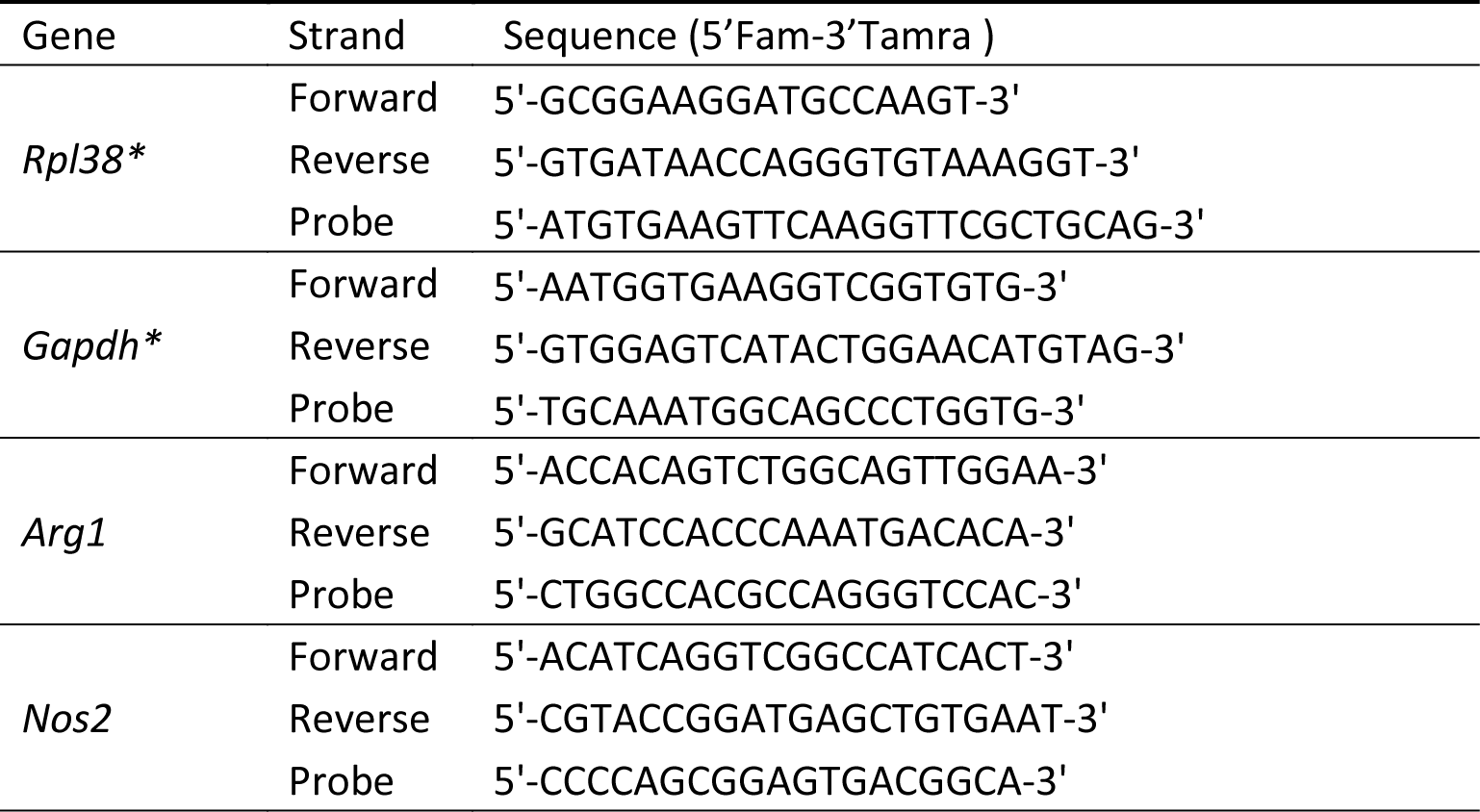
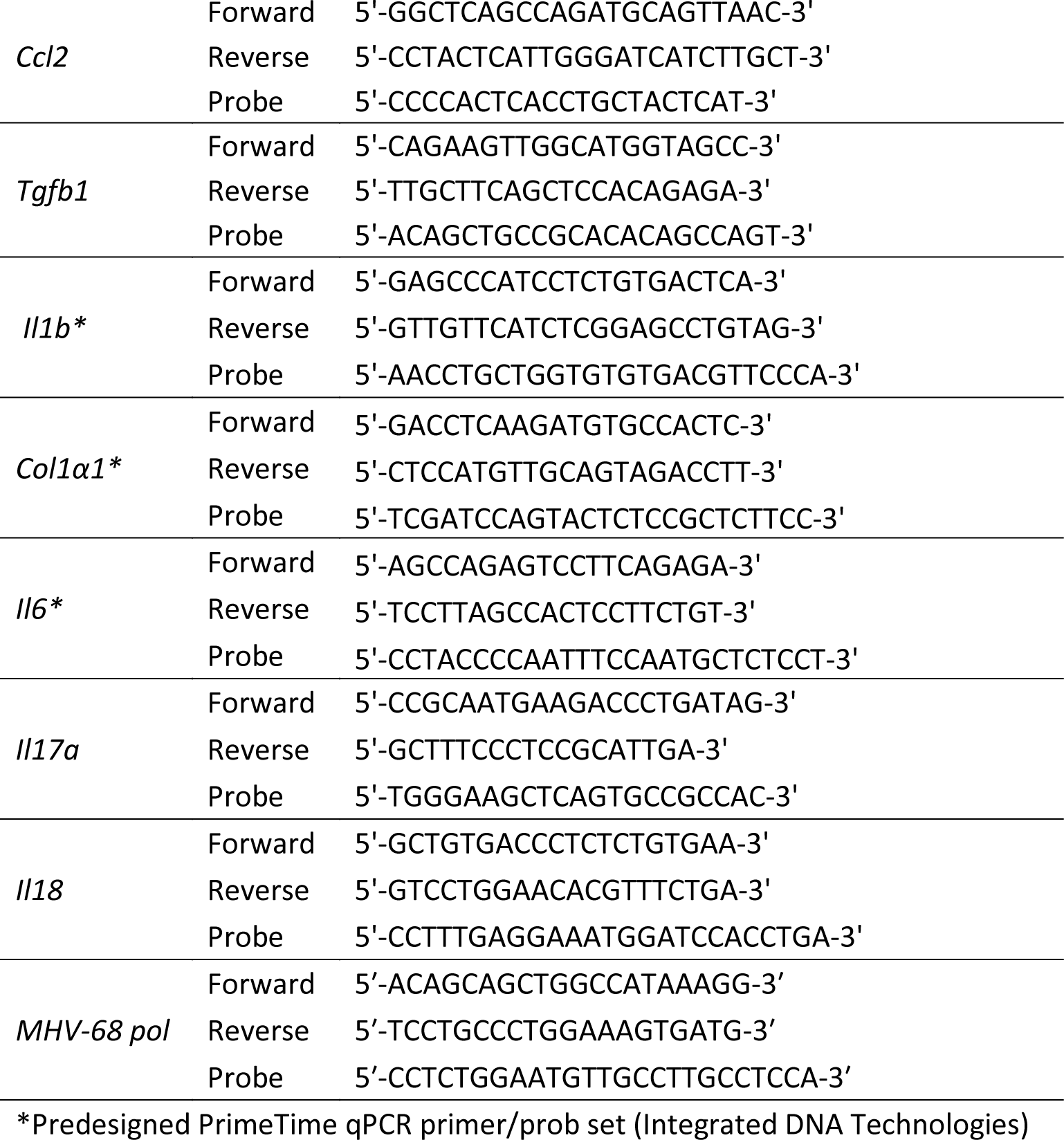
Primers and probes for quantitative PCR.

### Hydroxyproline Assay

The hydroxyproline content in lung tissue was quantified using an assay modified from previously described methods^32^. In summary, individual lung lobes were isolated and homogenized in 1 mL PBS and subsequently hydrolyzed through the addition of 1 mL of 12 N HCl (Sigma Aldrich) followed by incubation at 120°C for 18 hours. The assay was carried out in a 96-well plate, where 5 µl of each hydrolysate sample, as well as a series of standard concentrations of cis-4-Hydroxy-L-proline (Millipore Sigma), were incubated with a chloramine T (Millipore Sigma) solution for 20 minutes. This was followed by the addition of Ehrlich’s reagent (Millipore Sigma) and a further incubation at 65°C for 15 to 20 minutes to induce color development. The absorbance of each sample was then measured at 550 nm using a BioTek Synergy H1 Plate Reader (Agilent Technologies).

### Lung section and fibrosis scores

Whole lungs were prepared for fibrosis assessment by first being perfused and inflated before fixation in 10% buffered formalin. Following fixation, the tissues were subjected to dehydration through an ethanol series and embedded in paraffin. Tissue sections of 3 µm thickness were then sectioned, placed on glass slides, and stained using the Masson’s trichrome technique. Photomicrographs of the stained sections were captured using a Nikon Eclipse E400 microscope equipped with an Olympus EP50 camera. For each stained section, five fields were imaged at 100x magnification. The severity of fibrosis within these images was quantitatively evaluated using the Modified Ashcroft scoring method^82^.

### Lung cell preparation and flow cytometry analysis

To prepare single-cell suspensions from lung tissue for flow cytometry, whole lung collagenase digestion was performed as previously described^13^. Briefly, mice were euthanized with CO2, followed by perfusion of the lungs via the right ventricle with 5 mL of PBS. The lungs were then excised, and lung lobes were finely minced and incubated in 15 mL of complete media containing 1 mg/mL collagenase (Roche) and 17 U/mL DNase I (Sigma Aldrich) at 37°C for 30 minutes. Tissue homogenization was achieved by repeatedly drawing the digested mixture through the bore of a needleless 10-mL syringe, followed by filtration through a 100 μm mesh to obtain a suspension of single cells. For intracellular staining, cell suspensions were stimulated with 0.05 μg/mL phorbol 12-myristate 13-acetate (PMA) and 0.75 μg/mL ionomycin (both from Sigma Aldrich) in the presence of the GolgiStop protein transport inhibitor (BD Pharmingen) for 4 hours. Prior to staining, 1 x 10^6^ cells were blocked using anti-CD16/CD32 antibodies (Fc block; BD Pharmingen) and then labeled with a panel of fluorochrome-conjugated antibodies including CD45 (30-F11, BD Pharmingen), CD11c (N418, eBioscience), CD11b (M1/70, BD Horizon), I-A^b^ (AF6-120.1, BD Horizon), Siglec F (E50-2440, BD Horizon), CD103 (M290, BD Bioscience), Ly6G (1A8, BioLegend), CD90.2 (53-2.1, BD Pharmingen), PD-L1 (10F.9G2, BioLegend), Lg6C (HK1.4, BioLegend), Ep-CAM (G8.8, BioLegend), CD64 (X54-5/7.1, BioLegend), CD24 (M1/69, BioLegend), TCRγδ (GL3, BD Horizon), CD4 (GK1.5, BioLegend), PD-1 (J43, BD Horizon), CD3 (17A2, BD Pharmingen), CD8a (Ly-2, BioLegend), and CCR6 (29-2L17, BioLegend). After cell surface staining, the cells were then fixed and permeabilized with Foxp3 staining buffer set (eBioscience) and stained for intracellular TNFα (MP6-XT22, BioLegend), RORγT (Q31-378, BD Pharmingen), FoxP3 (FJK-16s, eBioscience), IFNγ (XMG1.2, BD Pharmingen), and IL-17a (TC11-18H10, BioLegend). Cell flow cytometry test was conducted on a Cytek Aurora analyzer (Cytek Biosciences). Data were analyzed by FlowJo software version 10.9.0 (FlowJo, LLC).

### Macrophage and CD4 T cell isolation

Macrophages were isolated from lung single-cell suspensions using microbeads conjugated to anti-F4/80 antibodies according to the manufacturer’s protocol (Miltenyi Biotec). CD4+ T cells were negatively selected from the lung single-cell suspensions utilizing the MojoSort™ Mouse CD4 T Cell Isolation Kit (BioLegend) in alignment with the manufacturer’s guidelines.

### Generation of bone marrow-derived dendritic cells (BMDCs)

BMDCs were generated from hematopoietic stem cells harvested aseptically from the femurs of 6-to 8-week-old C57BL/6J mice. The collected bone marrow cells were resuspended in RPMI 1640 medium (Gibco) supplemented with 10 ng/mL of recombinant murine GM-CSF (R&D Systems) to achieve a final concentration of 5 x 10^5^ cells/mL. One milliliter of the cell suspension was aliquoted into each well of a 24-well tissue culture plate (FisherBrand) and the cells were cultured for 6 days to facilitate differentiation into dendritic cells.

### Generation of induced TH17 (iTh17) cells

The *in vitro* differentiation of Th17 cells was conducted in accordance with a previously published protocol^83^. Briefly, splenocytes were harvested by mechanical disruption of the spleens from C57BL/6J or PD-1 knockout (PD-1KO) mice, passing the tissue through a 100 µm nylon cell strainer with the aid of a 3 mL syringe plunger. Following splenocyte collection, naïve CD4^+^ T cells were isolated using a negative selection kit (STEMCELL Technologies). The purified naïve CD4^+^ T cells, plated at a density of 3 x 10^5^ cells/well in a 96-well plate, were then stimulated with plate-bound anti-CD3 (clone 17A2, Invitrogen) and anti-CD28 (clone 37.51, Invitrogen) antibodies. Differentiation into Th17 cells was supported by a cytokine milieu consisting of recombinant IL-6, TGF-β, and IL-23 (all from R&D Systems), in conjunction with neutralizing antibodies against mouse IFNγ and IL-4 (also from R&D Systems). Culturing was performed in IMDM medium (Gibco) supplemented with the necessary growth factors. Cells were incubated for 3 days to promote the generation of induced iTh17 cells.

### Stimulation of BMDC with HK *L. johnsonii* and co-culture with iTh17 cells

BMDCs were incubated with HK *L. johnsonii* at a 1:2 ratio (BMDCs:HK *L. johnsonii*) at 37°C for 24 hours. Subsequently, the BMDCs were washed twice with PBS. To evaluate the surface expression of PD-L1 on BMDCs, aliquots of the cells were stained with antibodies against CD11c (clone N418, eBioscience), CD11b (clone M1/70, BD Horizon), I-Ab (clone AF6-120.1, BD Horizon), and PD-L1 (clone 10F.9G2, BioLegend).Stimulated BMDCs were then co-cultured with induced Th17 (iTh17) cells, which had been differentiated from naïve CD4+ T cells originating from either B6 or PD-1KO mice, using a BMDC:iTh17 cell ratio of 1:5. After 24 hours of co-culture, the concentration of IL-17A in the culture supernatant was quantified using the Mouse IL-17 DuoSet ELISA kit (R&D Systems) in accordance with the manufacturer’s directions.

### Determination of bacterial viability in lung homogenate

The lung homogenate was centrifuged at 300 x g for 2 minutes to pellet cell debris. The supernatant, typically 10 µL in volume, was then combined with an equal volume of 2X LIVE/DEAD *Bac*Light bacterial viability staining solution (Molecular Probes, Invitrogen). The mixture was kept in the dark for 15 minutes to allow the stain to penetrate the bacteria. The stained bacterial suspension was examined with a ZEISS Axioplan 2 fluorescence microscope to differentiate between live and dead bacteria. Images were captured using an Olympus DP28 camera to document and analyze the proportions of viable bacteria.

### Statistical analysis

Statistical analyses were conducted using GraphPad Prism version 9.0 (GraphPad Software). For comparisons between two groups, significance was assessed using 2-tailed Student’s t-tests for data with a normal distribution, and Mann-Whitney U tests were applied for data without a normal distribution. When comparing three or more groups, one-way ANOVA followed by Tukey’s multiple comparisons test was employed to evaluate the significance between groups.

## Notes

### Competing Interest Statement

The authors have declared no competing interest.

